# FusionESP: Improved enzyme-substrate pair prediction by fusing protein and chemical knowledge

**DOI:** 10.1101/2024.08.13.607829

**Authors:** Zhenjiao Du, Weimin Fu, Xiaolong Guo, Doina Caragea, Yonghui Li

## Abstract

To reduce the cost of experimental characterization of the potential substrates for enzymes, machine learning prediction model offers an alternative solution. Pretrained language models, as powerful approaches for protein and molecule representation, have been employed in the development of enzyme-substrate prediction models, achieving promising performance. In addition to continuing improvements in language models, effectively fusing encoders to handle multimodal prediction tasks is critical for further enhancing model performance using available representation methods. Here, we present FusionESP, a multimodal architecture that integrates protein and chemistry language models with a newly designed contrastive learning strategy for predicting enzyme-substrate pairs. Our best model achieved state-of-the-art performance with an accuracy of 94.77% on independent test data and exhibited better generalization capacity while requiring fewer computational resources and training data, compared to previous studies of finetuned encoder or employing more encoders. It also confirmed our hypothesis that embeddings of positive pairs are closer to each other in high-dimension space, while negative pairs exhibit the opposite trend. The proposed architecture is expected to be further applied to enhance performance in additional multimodality prediction tasks in biology. A user-friendly web server of FusionESP is established and freely accessible at https://rqkjkgpsyu.us-east-1.awsapprunner.com/.

## Introduction

Most enzymes are proteins capable of catalyzing a wide range of reactions within living organisms or under mild conditions *in vitro* with up to over a million-fold compared to spontaneous rates ^1,2^. Moreover, enzymes typically exhibit promiscuity, facilitating multiple reactions that may include physiologically irrelevant or potentially harmful processes ^3,4^. A comprehensive mapping of enzyme-substrate relationships will provide crucial guidance for future research in medicine, pharmaceuticals, bioengineering, and agriculture ^5–7^. However, it is prohibitively expensive to experimentally determine the catalytic interactions between molecules and enzymes. According to the UniProt Knowledgebase, approximately 10.8 million entries are related to enzymes, yet only 0.6% of these sequences have high-quality annotations of catalyzed reactions that are manually curated ^8^. There is a pressing need for high-throughput methods to address the scarcity of experimental validation in enzyme-substrate relationships.

Since the introduction of transformer architecture ^9^, various machine learning tasks have been revolutionized by leveraging pretrained large language models (LLMs) for molecular and protein representation, achieving significant advancements ^10–14,14,15^. Molecules and proteins can be represented as strings similar to natural language, using vocabularies such as the simplified molecular-input line-entry system (SMILES) for molecules and single-letter amino acid codes for proteins. Besides, protein and molecular databases (e.g., UniProt and ZINC) contained billions of protein sequences and molecular data, allowing to pretrain protein/chemistry LLMs. Based on their architecture, LLMs can be classified into encoder-only models (e.g., BERT - bidirectional encoder representations from transformers), decoder-only models (e.g., GPT - generative pre-trained transformers), and encoder-decoder models (e.g., T5 - text-to-text translation transformer). Among these, encoder-only architectures have gained popularity for remarkable performance in various downstream classification and regression tasks through transfer learning or fine-tuning ^5,10,12,14–16^. Recently, machine learning approaches for predicting compound-protein interaction (CPI) have increasingly adopted these encoder-only models ^17–20^, and enzyme-substrate pair prediction task is a subset of the CPI prediction task ^2,5,21,22^. Compared to traditional approaches using physicochemical parameters of molecules, enzyme sequences, or enzyme active site descriptors, employing pretrained BERT style encoders allows for better representation of proteins and molecules due to the domain knowledge learned via self-supervised learning from enormous datasets ^2,12,15,21–23^.

Enzyme substrate pairs prediction model is a multimodal prediction task, where two different “language” systems (i.e., amino acid sequences and SMILES) are involved during the embedding process. How to effectively leverage the enzyme embeddings and molecular embeddings generated from corresponding encoders becomes a crucial point in model performance enhancement. The most popular strategy is to simply concatenate the protein and molecule embeddings and this strategy and its modifications has been a common practice in the CPI discovery community ^18,20,24^. So far, the SOTA performance in enzyme-substrate pair prediction was achieved with the simple concatenation strategy in the study of Kroll et al., where a task-specific fine-tuned protein language model (PLM) (i.e., ESM-1b) and a pretrained domain-specific graph neural network (GNN) was used for embeddings and obtained an accuracy of 91.5% ^2^. Similar strategy was also used in the study of Song et al., where Kronecker products of protein and molecule embeddings as additional features were concatenated with the original embeddings to enhance CPI prediction performance ^19^. Beyond simple concatenation, cross-attention blocks have been employed to integrate local and high-level information from multiple protein and molecule encoders, capturing the relationship between protein and molecule representations for fused representation in CPI prediction ^17,18^. Recently, Kroll et al. introduced a multimodal BERT model for embedding protein-molecule complex, where the protein sequences and SMILES were input to a multimodal BERT model. They achieved SOTA performance across four datasets, including drug-target interactions, protein-small molecule interactions, enzyme-substrate Michaelis constants (KM), and substrate identification for enzymes, by concatenating PLM-generated protein embeddings, chemically learned molecule (CLM)-generated molecule embeddings, and multimodal encoder-generated protein-molecule embeddings ^5^. To maximize the prediction performance, these models either employed more encoders to enrich embedding protein and molecule or fused embeddings to enhance enzyme-substrate complex representations. However, these methods often require either complex fusion architecture design and substantial computational resources or additional datasets for finetuning, which impeded the deployment and application to different scenarios.

Inspired by the success of Contrastive Language-Image Pre-training (CLIP), where two independent encoders for images and texts were jointly trained to predict 400 million correct image-text pairs ^25^, we hypothesized that enzyme and substrate in a correct pair should be closer in a high dimensional space after projection, while unrelated components should be distinctly separated. This concept has been explored in single modality settings. To predict peptide-receptor complex pair, ESM-1b and ESM-MSA-1b were connected with independent encoder for peptide and protein receptor embeddings, respectively. These additional peptide and protein receptor encoders were jointly trained to capture inter-embedding similarities using a CLIP style model architecture ^26,27^. In contrast, Yu et al., (2023) took a different approach by designing a project head to refine embeddings without training additional encoders, thus reducing computation resource consumption. This strategy was employed in the single modality task of enzyme commission (EC) number prediction, where a frozen ESM-1b was used for enzyme embeddings, and a simple projection head was integrated with a contrastive loss function to refine embeddings ^28^. This approach achieved state-of-the-art (SOTA) performance across various benchmark datasets, particularly excelling in rare EC number predictions.

To the best of our knowledge, our model, FusionESP, is the first attempt to employ contrastive learning concept to address multimodality prediction tasks in biological science domain. Our objective was to develop an advanced machine learning module tailored for empowering existing LLMs to predict enzyme-substrate relationships across a wide range of proteins and molecules. This module aims to serve as a practical tool to streamline experimental processes and boost laboratory efficiency. In this study, we designed a simplified yet highly effective model architecture that employs a contrastive learning strategy. The architecture demonstrated remarkable performance enhancement in knowledge fusion within a multimodal context, even with limited data. Specifically, we utilized ESM-2 for enzyme embeddings and MolFormer for molecule embeddings. Rather than fine-tuning these encoders, adding additional encoders, or incorporating more features (e.g., extended-connectivity fingerprints), we designed projection layers for both encoders to refine and align the embeddings in the same high-dimensional space (Figure 1). Our model outperforms previous approaches with a simplified architecture and reduced computational demands during both training and inference phases.

**Figure 1.**
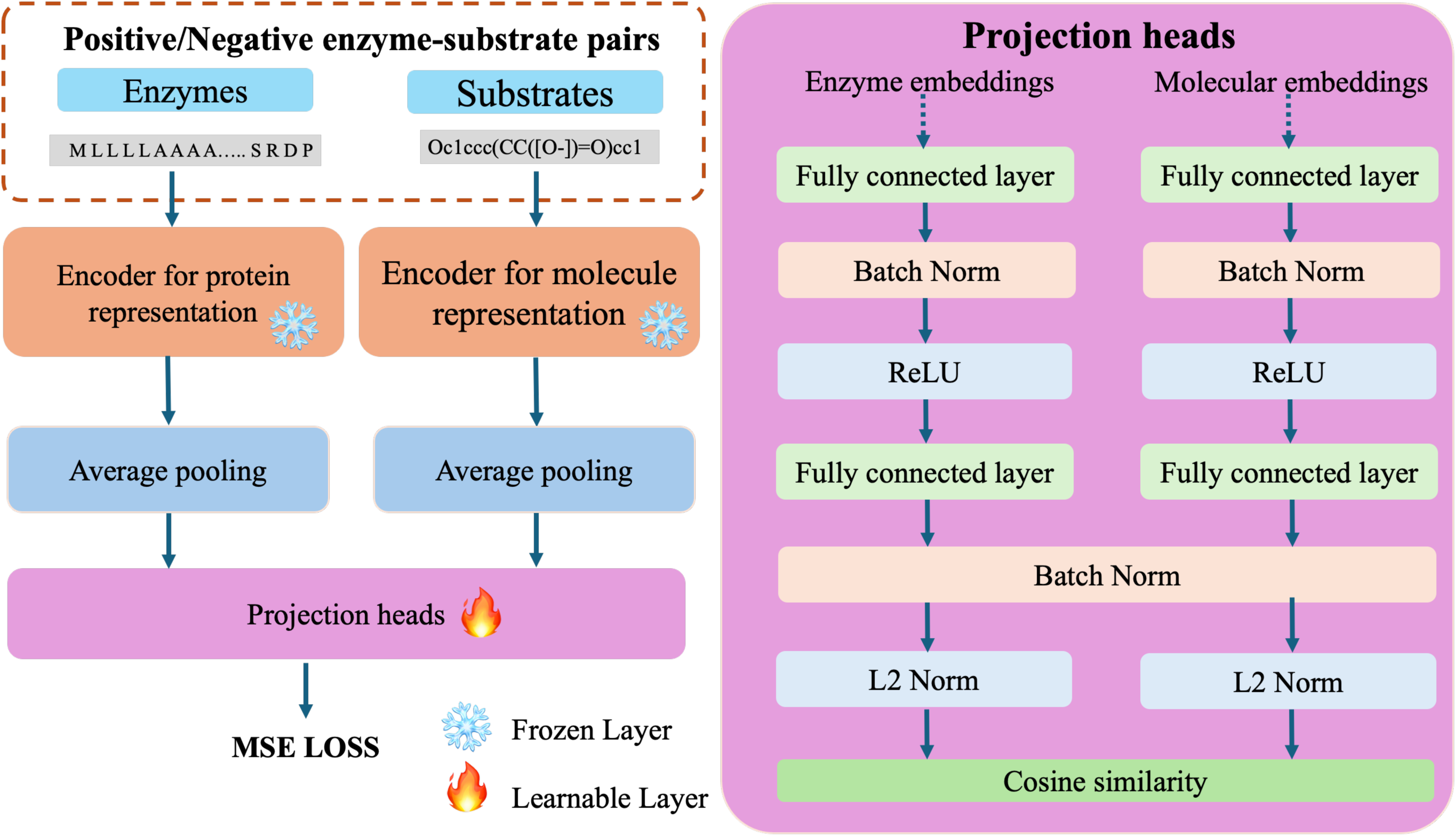
Model architecture for enzyme-substrate pair prediction model

## Methods

### Dataset

To ensure a fair comparison with previous studies, the datasets used in this study were sourced from existing studies ^2,5^. The positive enzyme-substrate pairs were from Gene Ontology (GO) annotation database, where entries have different levels of evidence: experimental, phylogenetically-inferred, computational analysis, author statement, curator statement, and electronic evidence. We utilized two datasets: one based on experimental evidence and the other on phylogenetic evidence. To challenge the model to distinguish true from false substrates, negative pairs were generated by randomly sampling three small molecules highly similar to the true substrates for the same enzyme sequences ^2^.

The experimental evidence-based dataset, originally split into training, validation, and test sets in the original study ^2,5^, contained 50,093 enzyme-substrate pairs in the training set, 5,422 pairs in the validation set, and 13,336 pairs in the test set. In contrast, the phylogenetic evidence-based dataset comprised a total of 765,635 pairs. These two datasets were downloaded from https://github.com/AlexanderKroll/ESP and https://github.com/AlexanderKroll/ProSmith, respectively.

Because of the computational demands of processing long protein sequences, we excluded some positive or negative enzyme-substrate pairs whose sequence embeddings exceeded hardware capacity (NVIDIA A100 GPU). Specifically, for ESM-2 models with output dimensions of 480 and 1280, sequences longer than 8000 were removed. Sixteen pairs were removed from the training dataset and no pair was removed from the test dataset. For the ESM-2 model with a 2560-dimensional output, sequences longer than 5500 were excluded. Correspondingly, 212 pairs were removed from the training dataset. Notably, no sequence in the validation and test datasets was removed.

### Calculating enzyme representation

In this study, enzymes were represented numerically using ESM2 models ^13,29^. ESM is a PLM project, initiated by Meta Fundamental AI Research (FAIR) in 2019 (https://github.com/facebookresearch/esm), which includes 19 pretrained PLMs with various dimensional output embeddings based on a modified Bidirectional Encoder Representations from Transformers (BERT) architecture. Those PLMs were trained on large protein sequence datasets (e.g., UR50/D 2021_04) through self-supervised learning. In this study, we employed three ESM-2 models, including the 480-dimensional output, 1280-dimensional output, and 2560-dimensional output, denoted as ESM-2-480, ESM-2-1280, and ESM-2-2560, respectively. The details about each PLM are listed in Supplementary Table S1. The amino acid one letter code of each enzyme was loaded into the PLM for embeddings. For an enzyme with *L* amino acids, the output of the ESM-2-*n* model for the enzymes is a *(L+1)*n* matrix, where *n* represents the output dimension of the ESM-2 model. The first row is the representation of the [sos] token, which is the indicator of “start of sentence” during training process. After that, each row in the matrix represented the corresponding amino acid from N-terminal to C-terminal with the consideration of the amino acid itself and the context of the entire sequence. We used an average pooling operation to unify the output dimension of enzymes with different lengths. Finally, any enzyme sequence loaded into the ESM-2-*n* for embeddings would result in a 1\**n* dimensional vector as its representation.

### Calculating molecule representation

MoLFormer was selected for numerical representation generation of molecules ^14^. MoLFormer was developed by IBM Research in 2022 and trained on canonical SMILES sequences of 1 billion molecules from ZINC database and 111 million molecules from PubChem with employment of rotary positional embeddings and linear attention mechanism. There was only one model available, called “MoLFormer-XL-both-10pct” trained with 10% of both datasets, at https://huggingface.co/ibm/MoLFormer-XL-both-10pct. Although the dataset size was smaller, this model demonstrated performance comparable to the full-size model. Therefore, we used this model for the molecule embeddings. We will use “MoLFormer” to refer this model in this paper. MoLFormer has a BERT-like architecture, similar to that of ESM models. To encode a canonical SMILES, the SMILES sequence was first converted into a numerical matrix, and then an average pooling operation was conducted to unify the feature dimension of SMILES sequences of different length into a 1*768 dimensional vector.

### Encoding models

ESM-2 models represent an upgrade over ESM-1b used in previous studies ^2,5^. They were trained on a new dataset (UR50/D 2021_04), while ESM-1b model was trained on UR50/S 2018_03 ^13,29^. Besides, the ESM-2 models employed rotary position embedding to replace the learned position embedding, which allowed it to embed any length of protein sequences, while ESM-1b can only embed sequences shorter than 1023 amino acids ^13,29^. As for MoLFormer, it was trained on a larger dataset than the one used in previous study (i.e., ChemBERTa-2) and employed more advanced algorithms (i.e., rotary position embedding and linear attention mechanism) ^5^. It exhibited better performance in one regression task (Lipophilicity dataset) and three classification tasks (BACE, BBBP, and ClinTox datasets) during downstream task performance evaluation ^10,14^.

### Model architecture design

The model architecture comprised two independent modules with similar inner architecture design for enzyme sequence input and molecule SMILES input, respectively (Figure 1). The enzyme module utilized an ESM model as the encoder for enzyme sequence embeddings, followed by average pooling to unify the embedding dimensions. These embeddings were then processed through a projection head to refine the dimension from 480/1280/2560 to 128. Similarly, the molecule module employed MoLFormer as the encoder, followed by average pooling and a projection head to refine the dimension from 768 to 128. Every enzyme substrate pair was finally transformed into two 128 dimensional vectors, where one represented the enzymes and the other represented the molecule. Cosine similarity between the two refined embeddings was calculated finally as the interaction portability of the pair. The value of the positive enzyme-substrate pairs was 1 and the true value of the negative pair was 0. A mean squared error (MSE) loss function was employed for the loss calculation as following:

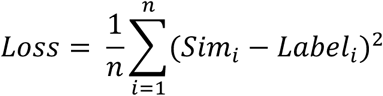

where n is the number of data points; *Sim_i_* is the cosine similarity of the 128 dimensional vectors of enzyme and substrate in the single pair; the *Label_i_* is the true label.

The projection head mainly referred to the projection heads in contrastive learning studies with a few modifications ^30–32^. The two projection heads for enzyme embeddings and molecular embeddings did not share weights and were trained simultaneously and independently through the MSE loss function except for the second batch norm layer. While the contrastive learning idea was inspired from CLIP ^25^, there was no huge dataset available for training, similar to the one that CLIP used to train the image and text encoders from scratch. Thus, in this study, we leveraged two pretrained encoders for embeddings to save the computational resources required for LLMs’ training and bypass the need for a huge enzyme-substrate pair dataset. At the same time, we employed the knowledge/expert-based negative dataset to address the challenges of scarcity in negative data points. The encoders were frozen and two independent projection heads were trained jointly as the learnable adaptor to maximize the model performance.

For each projection head, the number of neurons in the first fully connected layer corresponded to the encoder’s output dimension after average pooling. A batch normalization layer followed the fully connected layer before the ReLU activation function. Subsequently, the second fully connected layer (bottleneck layer) projected the input into a 128-dimensional vector, employing batch normalization and L2 normalization. The 128 dimension was selected mainly referring to SimCLR ^30^. The neurons in the first fully connected layer varied based on the encoder selected for model development. For instance, with the optimal combination of ESM-2-2560 and MoLFormer, the first fully connected layers had 2560 and 768 neurons for enzyme embeddings and molecule embeddings, respectively. Following projection into a 128-dimensional vector, both enzyme and molecule vectors shared the same batch normalization layer, assuming that they were at the same high-dimensional space.

### Model training

To expedite training and reduce computation needs, enzyme and molecule embeddings were pre-generated and saved for projection head training, bypassing the iterative generation of embeddings during training process. Adam optimizer was employed with default parameters. The batch size was 16 for the training on the experimental evidence-based datasets and 512 for the training on the phylogenetic evidence-based dataset. A relatively small batch size for experimental evidence-based datasets resulted from the small size of the dataset and the better capability of capturing the nuances, while a larger batch size was selected for the larger phylogenetic evidence-based dataset and captured the general features among positive/negative pairs.

The model trained on the experimental evidence-based dataset, denoted as FusionESP-exp, underwent 500 epochs, with the best-performing model checkpoint saved based on validation dataset performance. Training typically took 2-3 hours using a Tesla T4 GPU on Google Colab. The model trained on both the phylogenetic and experimental evidence-based datasets denoted FusionESP-XL, underwent similar processes, beginning with training on the phylogenetic evidence-based dataset for 500 epochs, followed by an additional 30 epochs on the experimental evidence-based dataset. The epoch numbers were selected based on learning curves (Supplementary Figure S1). The model, after training on a phylogenetic evidence-based dataset, was noted as FusionESP-phylo. At each state, best model checkpoint was saved based on validation dataset performance during this extended training process.

### Software

The neural network models were all implemented with Python and trained using the PyTorch library. The datasets and codes used to generate the results of this paper are available from https://github.com/dzjxzyd/FusionESP.

### Webserver

A user-friendly web server was deployed at Amazon Web Services (AWS) App Runner. The website was designed with html and css scripts, and the model deployment was achieved with Flask (2.2.2). Due to the constraint of computational resources, the FusionESP-XL with ESM-2-1280 and MolFormer was deployed for enzyme-substrate prediction. The web server supports large-scale processing, which allows users to upload their peptide information through xls or xlsx formats. The detailed scripts for web server development are available at https://github.com/dzjxzyd/FusionESP_server_1280.

## Results and Discussion

### Contrastive learning strategy exhibits superiority over simple concatenation strategy

The dataset was divided into training, validation, and test sets as in the original paper ^5^. Three different ESM-2 models and MoLFormer were employed as encoders for enzyme and molecule representations, respectively. FusionESP-exp was exclusively trained on the provided experimental evidence-based datasets. Although our training dataset was slightly smaller compared to those of previous models, all models, including ours, were evaluated on the same test dataset, ensuring a fair final performance comparison.

To assess the superiority of our contrastive learning strategy in enhancing model performance, we also implemented three counterpart baseline models with the popular concatenation strategy. We employed a simple two-layer neural network as the classifier to learn from the training data, details of which can be found in our GitHub repository. The results from all 6 models are shown in Table 1. All the models based on our contrastive learning strategy outperformed their counterparts, especially the FusionESP-exp with ESM-2-480 and MoLFormer, where the accuracy was 6.7% higher than that of the model based on simple concatenation strategy with ESM-2-480 and MoLFormer for embeddings. This underscores the significant advantage of our proposed contrastive learning strategy over the widely used concatenation strategy. A frozen BERT-based encoder paired with learnable neural networks as a projection head and contrastive loss function has previously shown success in single-modality conditions for EC number prediction ^28^. In this study, we employed two independent projection heads and an MSE loss function referring to CLIP model for multimodal prediction task. The promising results validated our hypothesis that positive enzyme-substrate pairs tend to be closer in high-dimensional space, while negative pairs are more distant. Moreover, due to the small size of the projection heads, our approach required a relatively modest dataset size, making it well-suited for biological applications with limited data availability.

**Table 1.**
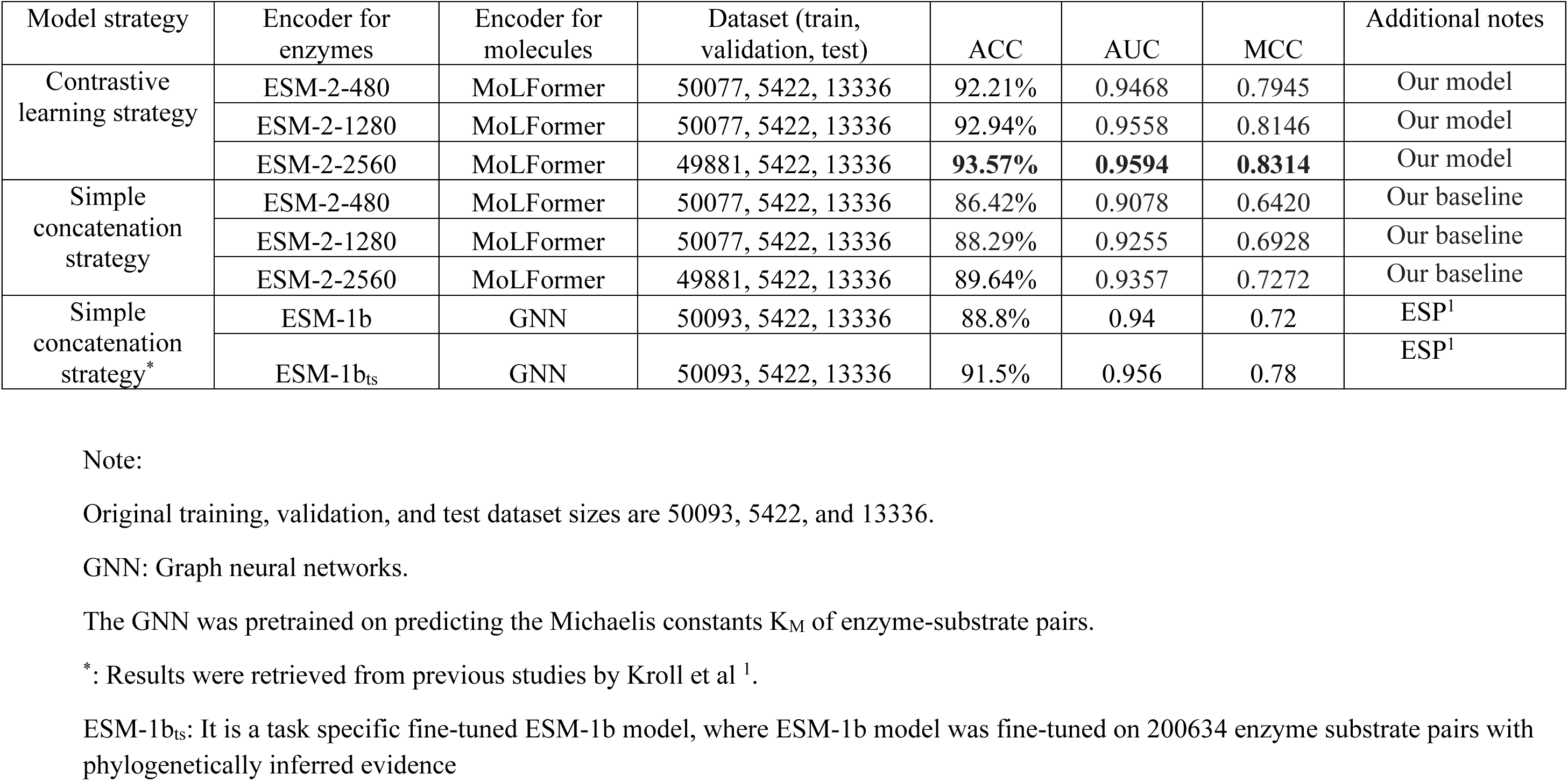
Performance of our model (FusionESP-exp) and the previous models trained on the experimental evidence-based dataset only.

| Model strategy | Encoder for enzymes | Encoder for molecules | Dataset (train, validation, test) | ACC | AUC | MCC | Additional notes |
| --- | --- | --- | --- | --- | --- | --- | --- |
| Contrastive learning strategy | ESM-2-480 | MoLFormer | 50077, 5422, 13336 | 92.21% | 0.9468 | 0.7945 | Our model |
|  | ESM-2-1280 | MoLFormer | 50077, 5422, 13336 | 92.94% | 0.9558 | 0.8146 | Our model |
|  | ESM-2-2560 | MoLFormer | 49881, 5422, 13336 | **93.57%** | **0.9594** | **0.8314** | Our model |
| Simple concatenation strategy | ESM-2-480 | MoLFormer | 50077, 5422, 13336 | 86.42% | 0.9078 | 0.6420 | Our baseline |
|  | ESM-2-1280 | MoLFormer | 50077, 5422, 13336 | 88.29% | 0.9255 | 0.6928 | Our baseline |
|  | ESM-2-2560 | MoLFormer | 49881, 5422, 13336 | 89.64% | 0.9357 | 0.7272 | Our baseline |
| Simple concatenation strategy* | ESM-1b | GNN | 50093, 5422, 13336 | 88.8% | 0.94 | 0.72 | ESP<sup>1</sup> |
|  | ESM-1b<sub>ts</sub> | GNN | 50093, 5422, 13336 | 91.5% | 0.956 | 0.78 | ESP<sup>1</sup> |

Note:
Original training, validation, and test dataset sizes are 50093, 5422, and 13336.
GNN: Graph neural networks.
The GNN was pretrained on predicting the Michaelis constants $K_M$ of enzyme-substrate pairs.
\*: Results were retrieved from previous studies by Kroll et al <sup>1</sup>.
ESM-1b<sub>ts</sub>: It is a task specific fine-tuned ESM-1b model, where ESM-1b model was fine-tuned on 200634 enzyme substrate pairs with phylogenetically inferred evidence

Additionally, a positive correlation between model performance and the size of ESM models/output dimensions was observed in both the contrastive learning and concatenation strategy groups. This trend aligns with findings from our previous studies ^33^, where larger model sizes/output dimensions encoded more information into the embeddings, resulting in improved overall performance.

### Model performance comparison with SOTA performance trained only on experimental evidence-based dataset

The FusionESP-exp models achieved superior performance compared to existing SOTA models, with accuracy, area under the curve, and Matthews correlation coefficient (MCC) ranging from 92.21% to 93.57%, 0.9468 to 0.9594, and 0.7945 to 0.8314, respectively. In Kroll et al.’s study, models combining ESM-1b or fine-tuned ESM-1b with a pretrained GNN on the same datasets outperformed models using ESM-1b and traditional extended-connectivity fingerprints (ECFP) for enzyme and substrate representation, which exhibited the superiority of pretrained model GNN over ECFP-based features ^2^. The performance of ESM-1b and GNN model achieved 88.8% accuracy, which was quite close to the one (ACC = 88.29%) of the FusionESP-exp composed of ESM-2-1280 & MoLFormer base models with simple concatenation strategy. The enzyme encoders in both models had the same model size (650 million parameters) and output dimension (1280 dimensions), with some modifications in ESM-2 regarding the training set and model architectures. As for the molecule encoders, MoLFormer was pretrained for general molecule representation, while GNN was pertained on a domain specific prediction task (i.e., production of Michaelis constants K_M_ of enzyme-substrate pairs) which was closer to our application scenario (i.e., enzyme-substrate pair prediction) ^2^. Besides, the pretrained GNN was still learnable during the training task on the enzyme-substrate pair dataset, and additional hyperparameter optimization for the gradient-boosting classifier was used to enhance the performance. Those reasons together contributed to slightly better performance of the ESM-1b & GNN based model over our simple concatenation models (ESM-2-1280 & MoLFormer), though we employed more advanced encoders for enzymes. When employing the contrastive learning strategy, FusionESP-exp with ESM-2-1280 & MoLFormer achieved 92.94% accuracy which was 5.27% higher than that of the model with ESM-1b & GNN. The second model from Kroll et al., employed a task specific fine-tuned ESM-1b model which was fine-tuned on 200,634 enzyme-substrate pairs with phylogenetically inferred evidence ^2^. The additional fine-tuning process enhanced the representation power of ESM-1b and increased the accuracy from 88.8 % to 91.5 %. However, compared to our models with contrastive learning strategy, the performance was inferior to ours by approximately 0.78-2.26 %, where no additional dataset, ESM-b fine-tuning and GNN fine-tuning were needed, highlighting the effectiveness of our proposed architectures for enzyme-substrate prediction tasks.

### Ablation study on the best-performed model

To further explore the function of each layer in the projection head, we conducted an ablation study on the best-performing FusionESP-exp model using ESM-2-2560 and MoLFormer with accuracy of 93.57% (as discussed in the previous section). We systematically investigated the effects of the first dense layer, ReLU activation function, bottleneck layer dimension, batch normalization layer, L2 normalization layer, and loss function. Table 2 summarized the results, indicating that the removal of either the first dense layer or entropy loss function had a detrimental effect on model performance. Given the original dataset’s 1:3 ratio of positive to negative pairs, these models were not trainable. Additionally, the bottleneck size, ReLU activation, and batch normalization layers contributed marginally to performance improvement, whereas the L2 normalization layer had negligible impact.

**Table 2.** Ablation study on FusionESP-exp with ESM-2-2560 and MoLFormer.

| Bottleneck size | First dense layer | ReLU | BatchNorm | L2 normalization | Loss function | ACC | AUC | MCC |
| --- | --- | --- | --- | --- | --- | --- | --- | --- |
| 32 | ✓ | ✓ | ✓ | ✓ | MSE | 93.09% | 0.9541 | 0.8192 |
| 64 | ✓ | ✓ | ✓ | ✓ | MSE | 93.02% | 0.9563 | 0.8177 |
| 128 | ✓ | ✓ | ✓ | ✓ | MSE | **93.57%** | **0.9594** | **0.8314** |
| 256 | ✓ | ✓ | ✓ | ✓ | MSE | 93.38% | 0.9574 | 0.8268 |
| 512 | ✓ | ✓ | ✓ | ✓ | MSE | 93.16% | 0.9562 | 0.8218 |
| 128 | × | ✓ | ✓ | ✓ | MSE | 74.31% | 0.8405 | 0.1310 |
| 128 | ✓ | × | ✓ | ✓ | MSE | 92.21% | 0.9499 | 0.7939 |
| 128 | ✓ | ✓ | × | ✓ | MSE | 92.01% | 0.9448 | 0.7901 |
| 128 | ✓ | ✓ | ✓ | × | MSE | 93.42% | 0.9591 | 0.8278 |
| 128 | ✓ | ✓ | × | × | MSE | 91.97% | 0.9447 | 0.7888 |
| 128 | × | × | ✓ | ✓ | MSE | 73.90% | 0.6867 | 0.0856 |
| 128 | ✓ | ✓ | ✓ | ✓ | CE* | 74.47% | 0.8217 | 0.4467 |

Note:
MSE: mean square error; CE: cross entropy
\* : to calculate the cross entropy, we employed the sigmoid function to get the probability based on similarity valued and then calculated the cross entropy loss between the probability and the label.

### Training the model with including phylogenetic evidence-based data further enhances its performance

In this section, we further enhanced model performance by training on a larger dataset combining phylogenetic evidence-based and experimental evidence-based datasets. Assuming the phylogenetic evidence-based dataset contained rich relationship information between enzymes and substrates, we directly trained models on this dataset for 500 epochs and subsequently the model underwent further training on the experimental evidence-based dataset for 30 epochs to optimize performance. Following the approach from a reference ^5^, we trained and evaluated the model performance on the same datasets for fair performance comparison. The results are shown in Table 3.

**Table 3.** Performance of our models (FusionESP-phylo and FusionESP-XL) and previous model trained on both experimental evidence-based and phylogenetic evidence-based datasets.

| Encoder for enzymes | Encoder for molecules | ACC | AUC | MCC | Additional notes |
| --- | --- | --- | --- | --- | --- |
| ESM-2-2560 | MoLFormer | 93.98% | 0.9567 | 0.8419 | Our model (FusionESP-phylo)* |
| ESM-2-2560 | MoLFormer | **94.77%** | 0.9653 | **0.8628** | Our model (FusionESP-XL)** |
| ESM-2-1280 | MoLFormer | 93.93% | 0.9534 | 0.8404 | Our model (FusionESP-phylo)* |
| ESM-2-1280 | MoLFormer | 94.56% | 0.9635 | 0.8572 | Our model (FusionESP-XL)** |
| ESM-1b | ChemBERTa-2 | 93.88% | 0.9517 | 0.8391 | Our model (FusionESP-phylo)* |
| ESM-1b | ChemBERTa-2 | 94.3% | 0.9618 | 0.8519 | Our model (FusionESP-XL)** |
| ESM-1b | ChemBERTa-2 | 94.2% | **0.972** | 0.85 | ProSmith<sup>2</sup> |

Note:
\* : the model (FusionESP-phylo) was trained on the phylogenetic evidence-based dataset only;
\*\* : the model (FusionESP-XL) was trained on the phylogenetic evidence-based dataset first and continued to be trained on the experimental evidence-based dataset, corresponding model.

Notably, all the models trained on phylogenetic evidence-based dataset only (FusionESP-phylo) outperformed all the models trained on experimental evidence-based dataset from Table 1 (FusionESP-exp), including models employing ESM-1b and ChemBERTa-2. The phylogenetic evidence-based dataset contained 764,449 data points, which was around 15 times larger of the experimental evidence-based dataset. We can infer that the number of learnable parameters was not yet saturated under current datasets and can be expected to further enhance performance with a larger dataset.

Similarly, larger encoder-based models exhibited better performance, with FusionESP-phylo and FusionESP-XL using ESM-2-2560, achieving accuracies of 93.98% and 94.77%, respectively, compared to 93.93% and 94.56% from models using ESM-2-1280. Besides, an improvement was also observed after the models were further trained on the experimental evidence-based dataset. This is not surprising, as the data points in training dataset from experimental evidence-based dataset were much closer to the data points in test dataset, and thus allowing the model to perform better in test dataset. There was also a performance improvement between the models with ESM-2-1280 and MoLFormer and the models with ESM-1b and ChemBERTa-2 under the same contrastive learning strategy. It mainly resulted from the two encoders. ESM-2-1280 and MoLFormer were trained with larger or updated datasets and advanced architectures (e.g., rotary position embeddings, linear attention mechanism) and achieved better performance than that of ESM-1b and ChemBERTa-2 in downstream prediction tasks ^10,13,14^. It is worth noting that ESM-1b can only embed enzyme sequences shorter than 1024 amino acids or truncated enzymes sequences with the first 1023 amino acids, which caused some information loss and inferior performance.

### The projection heads outperform an additional multimodal BERT

Compared to the previous SOTA model (ProSmith designed by Kroll et al.), our model trained and evaluated on the same datasets achieved superior performance ^5^. In the study of ProSmith, besides employing ESM1b and ChemBERTa-2 for enzyme and molecule embeddings, they introduced an additional multimodal BERT model trained on the combined text of the enzyme sequence and SMILES. The BERT model was pretrained on 1,039,565 data points from ligand target affinity dataset ^5^. Therefore, ProSmith employed three encoders for embeddings and connected them with an optimized gradient boost model for prediction, achieving an accuracy of 94.2% ^5^. In contrast, our model (FusionESP-XL) achieved 94.77% accuracy using only two pretrained encoders without additional pretraining of a new multimodal BERT model. With our contrastive learning strategy, FusionESP-XL with ESM-1b and ChemBERTa-2 only employed two encoders same as that of ProtSmith without the multimodal BERT achieved slightly higher accuracy.

In order to explore the contribution to the performance of our proposed architecture, we employed ESM1b and ChemBERTa-2 for enzyme and molecule embeddings and trained the model with our architecture with modifications according to the input dimensions. The performance was slightly better than that of ProSmith. Our promising performance was attributed to our simple projection heads, effectively fusing embeddings from two encoders for prediction tasks, marginally superior to the effect of an additional multimodal BERT encoder and well-optimized gradient boost classifier. At the same time, the ESM1b model was based on learned position embeddings, and it cannot embed the enzymes whose sequence length is longer than 1023 amino acids ^29^. Therefore, removed those enzyme-substrate pairs whose sequence length was longer than 1023. Though our training dataset was smaller than the one used in ProSmith, our performance was still comparable. Such a comparison further demonstrated the superiority of our proposed architecture.

### Evaluating FusionESP-XL performance in unseen enzymes and small molecules

In order to further characterize the model performance in rarely seen enzymes, we first split the test dataset into three subgroups: datapoints with enzymes with a maximal sequence identity to training data between 0 and 40%, between 40% and 60%, and between 60% and 80% based on CD-HIT ^34^. The results are presented in Table 4. It was observed that FusionESP-XL model with ESM-2-2560 & MolFormer achieved higher performance in enzyme-substrate pairs with higher sequence identity. Specifically, the accuracy, AUC, and MCC were 96.85%, 0.9871, 0.9198, respectively for enzymes with 60-80% sequence identity. The model still performed well for enzymes with 0-40% sequence identity with an accuracy of 93.07%, AUC of 0.9476, and MCC of 0.8185. It is worth noting that our model also exhibited better performance in the three subsets compared to the performance of ESP by approximately 1.95 - 4.57% in accuracy, respectively ^2^. Though trained on the same experimental evidence based dataset, FusionESP-exp with ESM-2-2560 & MolFormer still outperformed the ESP model across all three sub-datasets. When compared with ProtSmith, our model (FusionESP-XL with ESM-2-2560 & MolFormer) outperformed in most challenging sub-dataset (0-40 % maximal sequence identity) with the MCC increasing from 0.78 to 0.8185. This model also surpassed ProtSmith in the 40–60% maximal sequence identity sub-dataset in terms of MCC, while the MCC was slightly lower than ProtSmith in the 60–80% maximal sequence identity sub-dataset.

**Table 4.** The performance of FusionESP-XL and FusionESP-exp with ESM-2-2560 & MolFormer in rarely seen and unseen enzymes.

| Maximal sequence identity | ACC | AUC | MCC |
| --- | --- | --- | --- |
| **FusionESP-XL with ESM-2-2560 & MolFormer** |  |  |  |
| 0-40% | 93.07% | 0.9476 | 0.8185 |
| 40-60% | 96.12% | 0.9752 | 0.9001 |
| 60-80% | 96.85% | 0.9871 | 0.9198 |
| **FusionESP-exp with ESM-2-2560 & MolFormer** |  |  |  |
| 0-40% | 91.46% | 0.9334 | 0.7752 |
| 40-60% | 95.49% | 0.9731 | 0.8842 |
| 60-80% | 96.31% | 0.9864 | 0.9066 |
| ESP* |  |  |  |
| 0-40% | 89% | 0.93 | 0.72 |
| 40-60% | 93% | 0.97 | 0.83 |
| 60-80% | 95% | 0.99 | 0.88 |

Note: The performance of ProSmith was only exhibited in figure instead of table, and thus we did not have the exact values.
\*: The performance of ESP model was based on the task specific fine-tuned ESM-1b model.

Furthermore, we investigated the predictive capabilities of our models for small molecules with different frequency in training dataset (Table 5). Similar to that in maximal sequence identity, FusionESP-XL achieved better performance compared to FusionESP-exp across the 12 sub-datasets. Besides, the rarely seen small molecules tend to be wrongly predicted by the model, especially in unseen small molecules with a ACC of 80.93%. Compared to the performance in unseen small molecules in ProtSmith (MCC = 0.29) and ESP (MCC = 0.00), our model (FusionESP-XL with ESM-2-2560 & MolFormer) achieved slightly higher performance with a MCC of 0.2966. When it came to rarely seen small molecules (only present once in training dataset), our model’s performance was increased from 80.93 to 92.57%. Compared to ESP (MCC = 0.28) and ProtSmith (MCC= 0.69), our model achieved much higher performance (MCC = 0.7654).

**Table 5.** The performance of FusionESP-XL and FusionESP-exp with ESM-2-2560 & MolFormer in rarely seen and unseen small molecules.

| Number of positive samples with identical<br>metabolite in the training set | Size of Subset | ACC | AUC | MCC |
| --- | --- | --- | --- | --- |
| **FusionESP-XL with ESM-2-2560 & MolFormer** |  |  |  |  |
| 0 | 640 | 80.93% | 0.6942 | 0.2966 |
| 1 | 498 | 92.57% | 0.9445 | 0.7654 |
| 2 | 431 | 93.27% | 0.8935 | 0.7880 |
| 3 | 586 | 96.07% | 0.9657 | 0.8758 |
| 4 | 351 | 95.44% | 0.9634 | 0.8631 |
| 5 | 414 | 93.48% | 0.9486 | 0.7976 |
| 6 | 278 | 94.96% | 0.9597 | 0.8547 |
| 7 | 313 | 94.57% | 0.9731 | 0.8460 |
| 8 | 222 | 92.79% | 0.9724 | 0.7938 |
| 9 | 317 | 94.32% | 0.9575 | 0.8471 |
| 10 | 157 | 96.18% | 0.9900 | 0.8954 |
| >10 | 8876 | 95.86% | 0.9783 | 0.8995 |
| **FusionESP-exp with ESM-2-2560 & MolFormer** |  |  |  |  |
| 0 | 640 | 76.72% | 0.6637 | 0.1343 |
| 1 | 498 | 87.95% | 0.8870 | 0.6198 |
| 2 | 431 | 88.86% | 0.8738 | 0.6503 |
| 3 | 586 | 93.86% | 0.9584 | 0.8080 |
| 4 | 351 | 92.02% | 0.9397 | 0.7660 |
| 5 | 414 | 92.03% | 0.9432 | 0.7504 |
| 6 | 278 | 92.45% | 0.9580 | 0.7833 |
| 7 | 313 | 94.89% | 0.9625 | 0.8543 |
| 8 | 222 | 91.44% | 0.9432 | 0.7512 |
| 9 | 317 | 94.01% | 0.9704 | 0.8371 |
| 10 | 157 | 95.54% | 0.9737 | 0.8775 |
| >10 | 8876 | 95.30% | 0.9733 | 0.8855 |

The performance of our model in rarely seen and unseen enzyme and small molecules indicated that our model has better generalization capability. The projection heads can not only contribute to the increase in overall performance, but also enable the model to perform well in unseen data points.

To further explore the model performance under different maximal sequence identity and frequency of small molecules in training dataset, we further explore the model’s performance under different frequencies of small molecules in each sequence similarity subgroups (Figure 2). It was observed that within the same maximal sequence identity subgroup, performance improved as the frequency of small molecules in the training dataset increased. For example, in the 0–40% maximal sequence identity subgroup, the accuracy increased from 76.13% for unseen small molecules to 90.72% for those with a frequency ranging from 1 to 10, and further to 94.26% for frequencies greater than 10. Similarly, the prediction performance for unseen and rarely seen small molecules also improved as the maximal sequence identity increased. Particularly, the accuracy for those unseen small molecules was increased from 76.13% with 0 – 40 % maximal sequence identity to 85.09% for enzymes with 40 – 60 % maximal sequence identity and 93.07% for enzymes with 60 – 80 % maximal sequence identity. Predicting enzyme-substrate pairs was particularly challenging when the enzyme exhibited 0 – 40% sequence identity and the small molecule was unseen. However, when the enzyme had a higher maximal sequence identity, or the small molecules were present in the training dataset the model was able to make significantly more reliable predictions.

**Figure 2.**
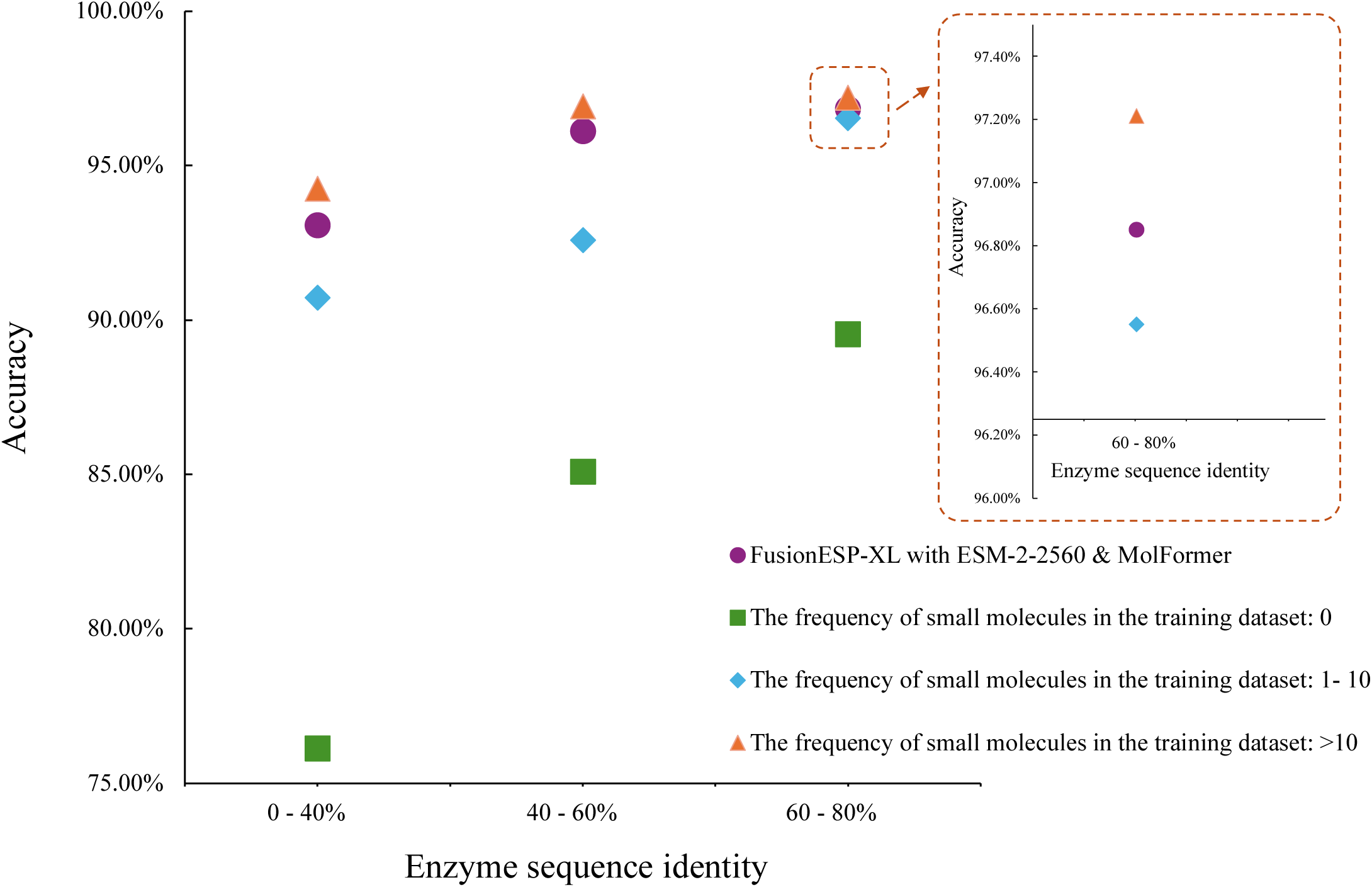
Prediction performance of FusionESP-XL with ESM-2-2560 & MolFormer. Note: We divided the test dataset into subset with different level of enzyme sequence identity and frequency of small molecules in the training dataset. Source data are provided as a Source Data file.

### FusionESP-XL models can express uncertainty

Internally, our model can also provide the cosine similarity score instead of only the positive or negative prediction results to interpret how confident the model is regarding its prediction. In this paper, we set 0.5 as the threshold for output results, where a cosine similarity score between an enzyme and a small molecule was predicted as a positive pair if the score is higher than 0.5, and otherwise it is a negative pair. The cosine similarity score can be also provided as output at our webserver at https://78k6imn5wp.us-east-1.awsapprunner.com for single reaction prediction or large-scale prediction.

To provide a more detailed assessment of prediction accuracies, Figure 3 displays the distributions of true (blue) and false (orange) predictions within our test dataset across various prediction scores. Most correct predictions had scores either close to 0 or close to 1, indicating that FusionESP-XL made predictions with high confidence. In contrast, false predictions were distributed more evenly across the prediction score range. The predictions for the data points with scores between 0.4 and 0.6 were more likely to be incorrectly predicted. Therefore, for practical usage of FusionESP-XL, input pairs with cosine similarity score within the 0.4 to 0.6 range should be treated as uncertain and used cautiously in decision making.

**Figure 3.**
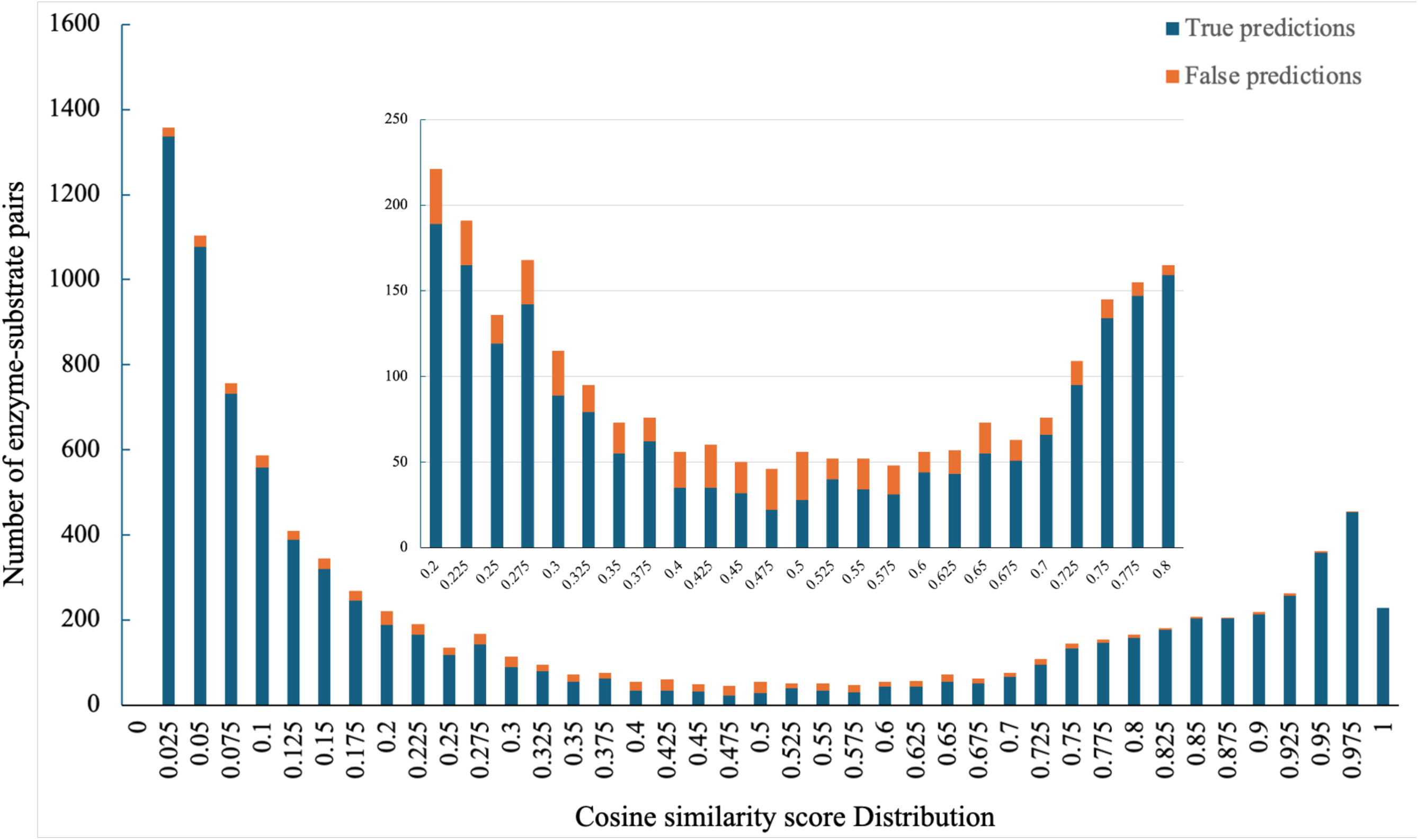
Prediction scores around 0.5 indicate model uncertainty. Note: Stacked histogram bars display the prediction score distributions of true predictions (blue) and false predictions (red). The inset shows a blow-up of the interval [0.2, 0.8]. Scores are prediction by FusionESP -XL with ESM-2-2560 and MoLFormer. Source data are provided as a Source Data file.

## Conclusion

Overall, our proposed contrastive learning strategy (FusionESP) achieved SOTA performance, leveraging two frozen encoders and two simple projection heads with an MSE loss function. The best model, FusionESP-XL, which utilized ESM-2-2560 and MoLFormer, achieved SOTA performance with accuracy, AUC, and MCC of 94.77%, 0.9653 and 0.8628, respectively. Even under limited data availability (using an experimental evidence-based dataset only), our model FusionESP-exp with ESM-2-2560 and MoLFormer also achieved SOTA performance with accuracy, AUC, and MCC of 93.57%, 0.9594 and 0.8314, respectively. Furthermore, our models can handle proteins of any length, unlike ProSmith, which truncates all enzymes into 1023 amino acids due to constraints from the ESM-1b models. This truncation can lead to confusion when two different enzymes share the same first 1023 residues from N- to C-terminus. Moreover, our strategy does not require extra pretrained encoders or datasets for pretraining, but exhibited better generalization capability, whereas ProSmith’s development relied on a highly correlated dataset (ligand-target affinity) to pre-train the multimodal BERT encoders before applying them to enzyme-substrate complex embedding. Our model (FusionESP-XL with ESM-2-2560 and MoLFormer) shows great potential for future enzyme-substrate pair prediction, mapping reliable relationship between enzymes and their substrates. The contrastive learning strategy proposed in this study holds significant potential for other multimodal prediction tasks.

## Acknowledgements

This is contribution No. 25-059-J from the Kansas Agricultural Experimental Station. This work was supported in part by the National Science Foundation (2419880) and the Plant Protein Innovation Center.

## Data Availability

All source codes used in this publication are freely available for academic use under an MIT license at https://github.com/dzjxzyd/FusionESP. The datasets can be download at https://zenodo.org/records/13891018.

## Supporting Information

Figures S1 shows learning curves of the model on the phylogenetic evidence-based dataset (a) and the experimental evidence-based dataset (b). Table S1 shows pretrained protein language model in this study.

---

**Table S1.**
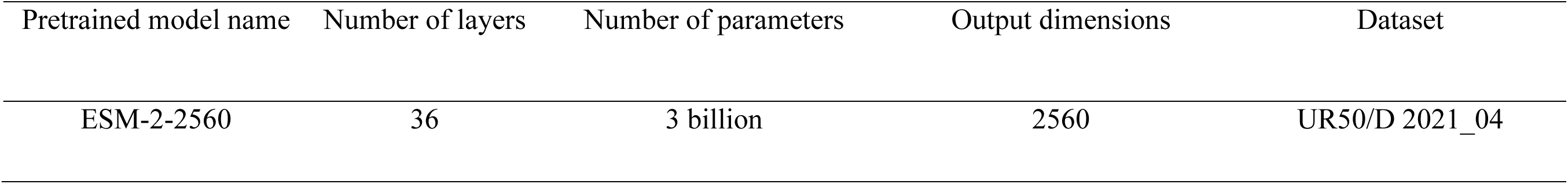
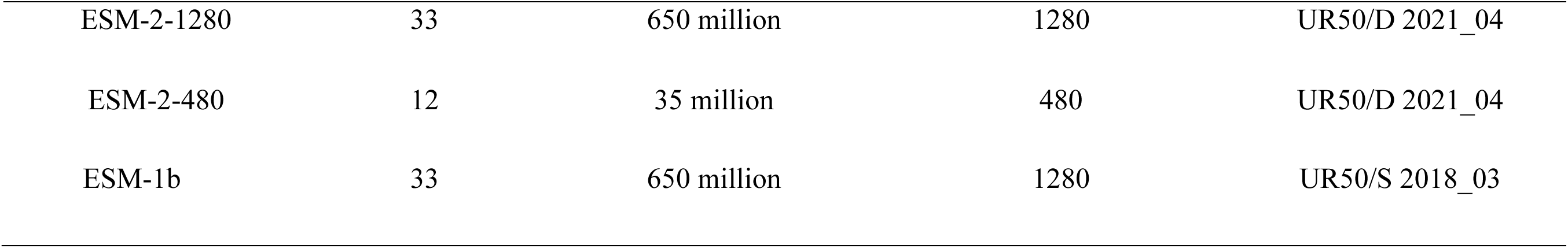
Pretrained protein language model in this study.

| Pretrained model name | Number of layers | Number of parameters | Output dimensions | Dataset |
| --- | --- | --- | --- | --- |
| ESM-2-2560 | 36 | 3 billion | 2560 | UR50/D 2021_04 |
| ESM-2-1280 | 33 | 650 million | 1280 | UR50/D 2021_04 |
| ESM-2-480 | 12 | 35 million | 480 | UR50/D 2021_04 |
| ESM-1b | 33 | 650 million | 1280 | UR50/S 2018_03 |

**Table S1** Pretrained protein language model in this study
| Pretrained model name | Number of layers | Number of parameters | Output dimensions | Dataset |
| --- | --- | --- | --- | --- |
| ESM-2-2560 | 36 | 3 billion | 2560 | UR50/D 2021_04 |
| ESM-2-1280 | 33 | 650 million | 1280 | UR50/D 2021_04 |
| ESM-2-480 | 12 | 35 million | 480 | UR50/D 2021_04 |
| ESM-1b | 33 | 650 million | 1280 | UR50/S 2018_03 |

**Figure S1.**
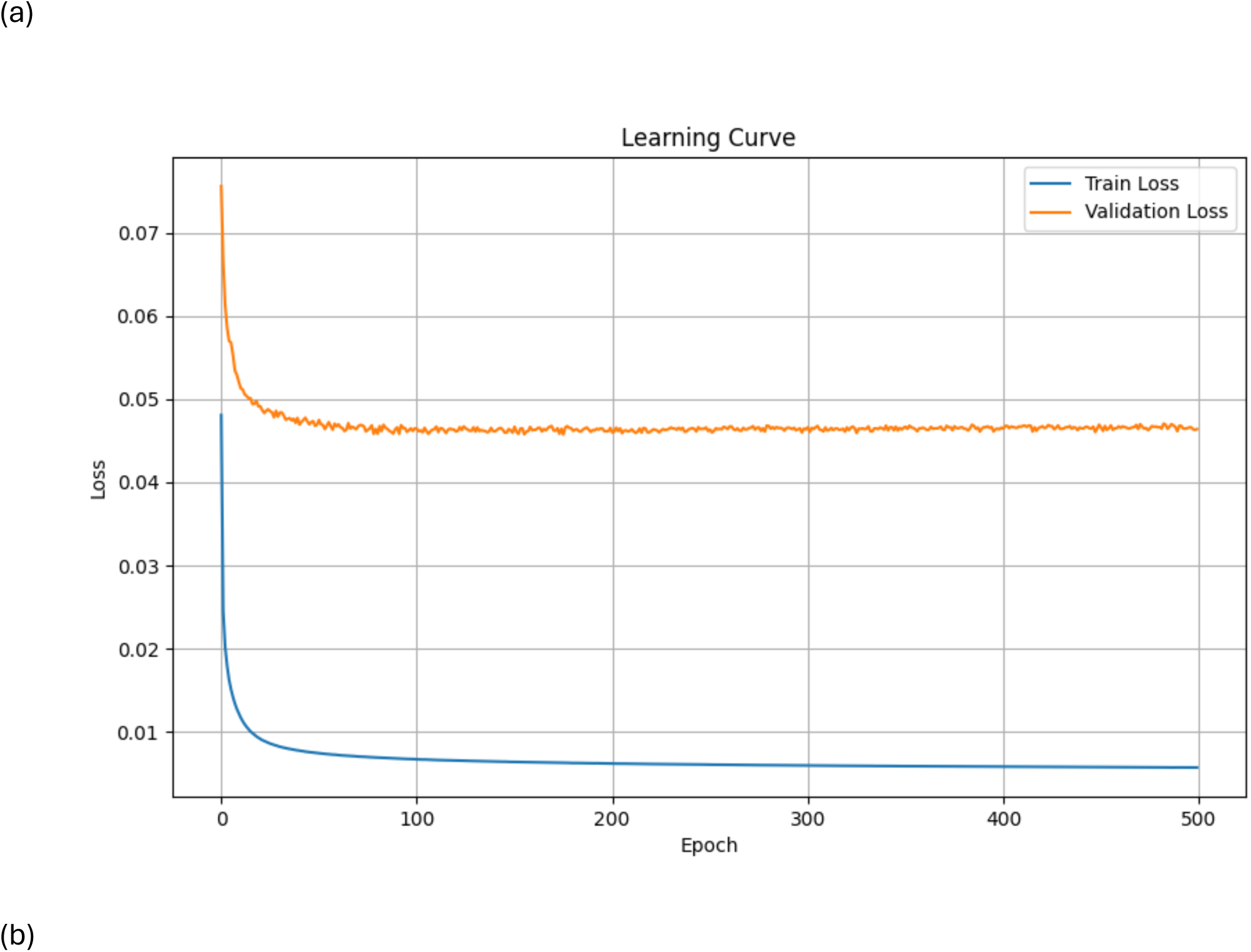
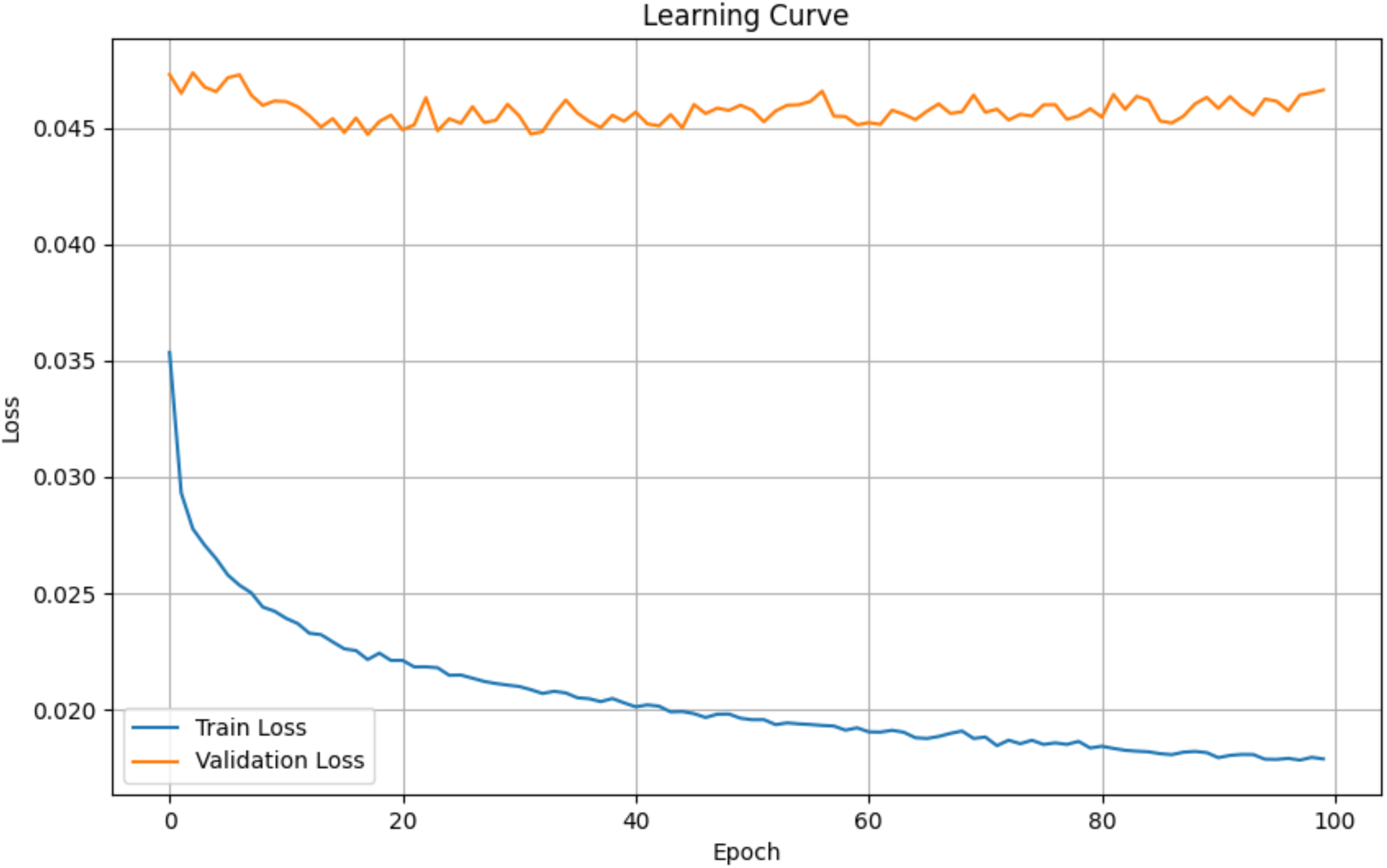
Learning curves of the model on the phylogenetic evidence-based dataset (a) and the experimental evidence-based dataset (b)

